# Validity Assessment of Michigan’s Proposed qPCR Threshold Value for Rapid Water-Quality Monitoring of *E. coli* Contamination

**DOI:** 10.1101/2022.07.16.500300

**Authors:** James N. McNair, Molly J. Lane, John J. Hart, Alexis M. Porter, Shannon Briggs, Benjamin Southwell, Tami Sivy, David C. Szlag, Brian T. Scull, Schuyler Pike, Erin Dreelin, Chris Vernier, Bonnie Carter, Josh Sharp, Penny Nowlin, Richard R. Rediske

## Abstract

Michigan’s water-quality standards specify that *E. coli* concentrations at bathing beaches must not exceed 300 *E. coli* per 100 mL, as determined by the geometric mean of culture-based concentrations in three or more representative samples from a given beach on a given day. Culture-based analyses require 18–24 h to complete, so results are not available for issuing beach notifications (advisories or closings) until the day following collection. This one-day delay is problematic because *E. coli* concentrations at beaches can change markedly from one day to the next. qPCR-based *E. coli* concentrations, by contrast, can be obtained in only 3–4 h, making same-day beach notifications possible. Michigan has proposed a qPCR threshold value (qTV) for *E. coli* of 1.863 log_10_ gene copies per reaction as a potential equivalent value to the state standard, based on statistical analyses of a set of training data from 2016–2018. The main purpose of the present study is to assess the validity of the proposed qTV by determining whether the implied qPCR-based beach notification decisions agree well with culture-based decisions on two sets of test data (from 2016–2018 and 2019–2020), and whether performance of the proposed threshold is similar on the test and training data. The results show that performance of the proposed qTV on both sets of test data was consistently good (e.g., 95% agreement with culture-based beach notification decisions during 2019–2020) and was at least as good as its performance on the training data set, supporting its use as an equivalent value to the state standard.

## 1 Introduction

Epidemiological studies have shown that total-body contact with recreational waters contaminated by fecal indicator bacteria (FIB) is associated with increased incidence of gastrointestinal illness (Stevenson, 1953; Dufour, 1984; Wade et al., 2008; Colford et al., 2012; EPA, 2012). Beaches of many recreational waterbodies in the United States are therefore monitored for FIB such as *Escherichia coli* (*E. coli*) and enterococci to assess the need for issuing beach notifications (advisories or closings) to protect human health. Numerical water-quality standards (WQS) for making beach notification decisions are set by states at FIB concentrations thought to provide adequate protection, based on the epidemiological studies just mentioned and guidance by the US Environmental Protection Agency (EPA) (EPA, 2012).

Michigan has approximately 1,390 identified public beaches, of which about 450 are monitored for *E. coli* (Michigan EGLE, 2018). Monitoring of beaches in Michigan is voluntary and is conducted by local health departments, which are required to comply with state WQS for total-body-contact recreation (Appendix A, Section A.1). These standards specify that water at beaches must not contain more than 300 *E. coli* per 100 mL, based on the geometric mean of culture-based concentrations in three or more representative samples from a given beach on a given day. Culture-based analyses, however, require 18–24 h to complete, so beach notification decisions cannot be made until the day after samples are collected. This one-day delay is problematic because beach FIB concentrations can change markedly from one day to the next (Whitman et al., 1999; Boehm et al., 2002; Whitman and Nevers, 2004; Boehm, 2007). EPA Draft Method C was developed to address this issue (Sivaganesan et al., 2019; Aw et al., 2019; Lane et al., 2020a). It is based on the quantitative real-time polymerase chain reaction (qPCR) and produces results in only 3–4 h, so same-day beach notification decisions would be possible if an appropriate qPCR threshold value (qTV) were established.

Toward this end, Michigan’s Department of Environment, Great Lakes, and Energy has proposed a qTV of 1.863 log_10_ gene copies per reaction (GC/reaction) for beach notifications. This threshold is based on a study by EPA researchers that analyzed the statistical relationship between culture-based and qPCR-based *E. coli* concentrations in a subset of samples collected as part of Michigan’s annual beach monitoring program for 2016–2018 (Haugland et al., 2021). To date, however, the proposed qTV has not been validated by assessing its performance on other data.

Michigan was the first state to implement qPCR-based beach monitoring on a state-wide basis. In 2016, the state’s beach monitoring program began determining culture-based and qPCR-based *E. coli* concentrations in split samples from more than 100 freshwater recreational (bathing) beaches across the state. The beaches represent diverse waterbodies that include three Laurentian Great Lakes, numerous inland lakes, and several rivers and streams. Each sample is split into two subsamples, one of which is analyzed using the culture-based IDEXX Colilert-18^®^ method (IDEXX Laboratories, Westbrook, ME) and the other using qPCR-based EPA Draft Method C. The resulting set of bivariate data permits characterization of the statistical relationship between the two methods of determining *E. coli* concentrations, as well as assessment of the ability of the proposed qTV to correctly predict culture-based beach notification decisions.

The purpose of the present study is to assess the validity of Michigan’s proposed qTV by characterizing and comparing its performance on beach monitoring data for 2016–2018 (6,564 samples) and 2019–2020 (3,205 samples), and to compare the numerical values of three key measures of the decision threshold’s performance calculated for the data used to derive the proposed qTV (the “training” data) with values of the same three performance measures calculated for our 2016–2018 and 2019–2020 data sets (the “test” data). Our 2016–2018 data set includes some of the data used in deriving the proposed threshold but contains many other data, as well (see Section 2.6). The 2019–2020 data apply to a different time period and were not involved in developing the threshold, so they permit a fully independent assessment of its validity.

Our assessment has two parts. First, we visually compare general properties of the two test data sets, including the overall distributions of the data, the distributions of the two subsets of each data set that do and do not exceed Michigan’s culture-based standard for beach notifications, and two overall measures of the degree to which qPCR-based estimates of *E. coli* concentration permit one to discern the difference between samples that do and do not exceed the culture-based standard (Section 3.1). Second, we compare performance of the proposed qTV quantitatively on the 2016–2018 and 2019–2020 test data, and on the 2016–2018 training data, using several numerical performance measures commonly employed for assessing decision or classification rules in various disciplines (Section 3.2).

## 2 Material and methods

Our assessment is based on existing data produced by multiple labs for Michigan’s beach monitoring program during 2016–2020. Sampling and analytical methods employed by contributing labs are detailed by Lane et al. (2020a,b) and outlined in this section and Appendix A. This section also identifies known limitations of the data and describes data-analysis methods used in the assessment.

### 2.1 Sampling locations

Data used in the assessment are derived from samples collected from over 100 bathing beaches on coastal and inland waters of Michigan (Fig. 1).

**Figure 1.**
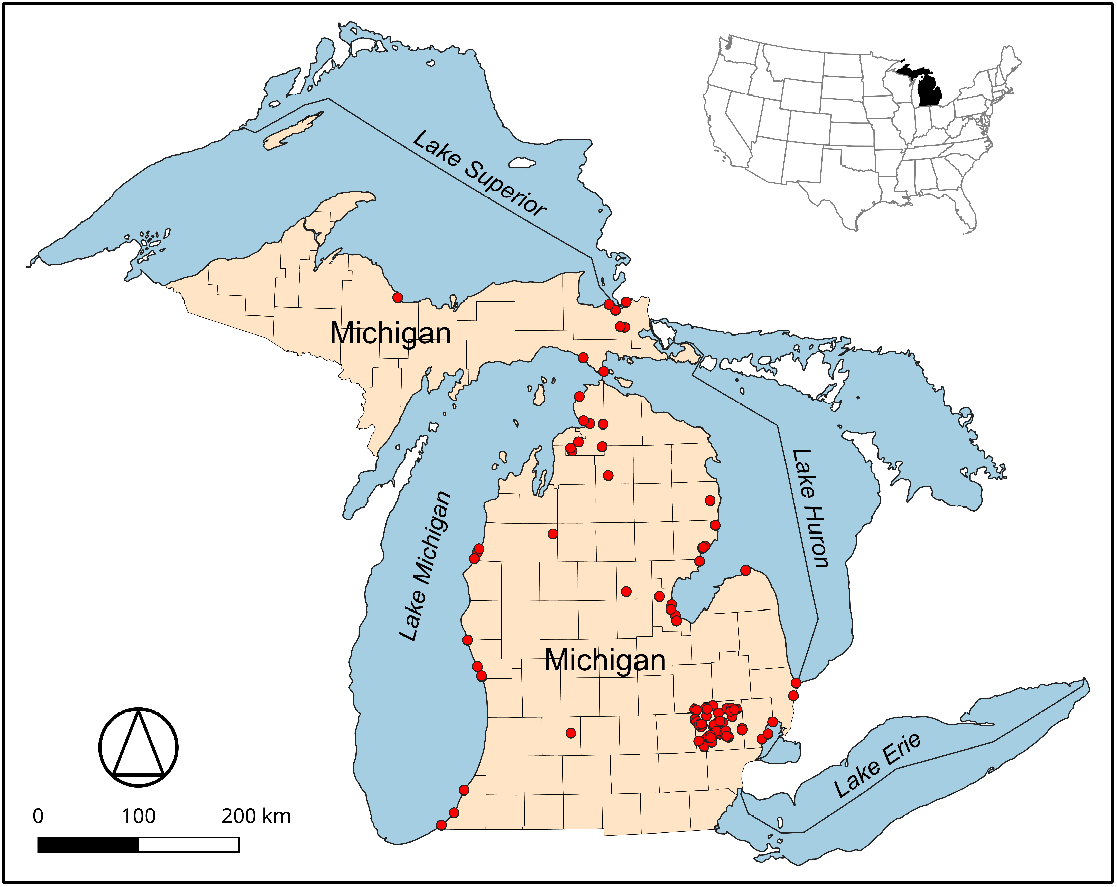
Locations of Michigan recreational beaches (dots) from which paired culture-based and qPCR-based data for 2016–2018 and 2019–2020 used in the present assessment were derived. Base map: Michigan Geographic Framework.

### 2.2 Contributing laboratories

The 2016–2018 data were contributed by ten Michigan labs, the 2019–2020 data by seven (Appendix A, Table A.1). All labs performed culture-based analyses using Colilert-18 (Section 2.4) and qPCR analyses using EPA Draft Method C (Section 2.5).

### 2.3 Sample collection and preparation

Sampling methods are described in detail by Lane et al. (2020a). Briefly, beaches were sampled from May through October each year at frequencies that varied according to the sampling plan of the local health department. During each sampling event, separate water samples (typically 200–500 mL) were collected in sterile bottles at 3–6 representative locations, depending on beach size. Samples were stored at 4 °C for transport and processed within 6 h of collection. Samples from each beach and day were either processed and analyzed individually or were analyzed as a single composite, according to the local sampling plan. The *E. coli* concentration in each individual or composite sample was determined using culture-based and qPCR methods described in Sections 2.4 and 2.5.

### 2.4 Colilert-18 *E. coli* quantification

The most probable number (MPN) of *E. coli* colony-forming units per 100 mL (MPN/100 mL) was determined for each sample using the Colilert-18 Quanti-Tray/2000^®^ IDEXX system following manufacturer instructions. The lower limit of quantification (LLOQ) was 1.0 MPN/100 mL; the upper limit of quantification (ULOQ) was 2419.6 MPN/100 mL.

### 2.5 qPCR *E. coli* quantification

qPCR-based *E. coli* concentrations as log_10_ GC/reaction were determined using EPA Draft Method C (Sivaganesan et al., 2019; Aw et al., 2019; Lane et al., 2020a). The qPCR assays EC23S857 *E. coli* (Chern et al., 2011) and Sketa22 salmon DNA (EPA, 2012), method of DNA extraction, and method of qPCR analysis employed by the contributing labs are detailed by Lane et al. (2020a,b) and summarized in Appendix A, Section A.3. Standard curves were constructed as described by Lane et al. (2020b), using standards prepared and quantified by EPA’s Office of Research and Development (ORD) lab in Cincinnati, Ohio, as described by Sivaganesan et al. (2019). LLOQs were determined separately for each standard curve, as described by Lane et al. (2020b).

### 2.6 Known data limitations

As noted above, data used in the present assessment were produced by multiple labs for Michigan’s annual beach monitoring program during 2016–2020 and thus were not produced specifically for this assessment. As is typical of large environmental data sets comprising data from multiple labs and years, the beach monitoring data have several limitations, and it is important to be aware of these when assessing validity of the proposed qTV.

First, though Michigan recreational WQS prescribe separate analysis of three or more discrete samples from each beach, local health departments sometimes composite the samples before analysis as a cost-saving measure to permit monitoring more beaches. In such cases, the raw data for a sampling event consist of a single qPCR-based concentration and a single culture-based concentration instead of multiple concentrations of each type.

A second limitation is that roughly half of the samples (53%) yielded qPCR-based or cultured-based (or both) *E. coli* concentrations that are censored (i.e., outside the range of quantification). Specifically, many samples had qPCR-based concentrations below the LLOQ, and many had culture-based estimates either below the LLOQ or, less commonly, above the ULOQ for Colilert-18. (No sampling event produced samples for a given beach and date that included both <LLOQ and >ULOQ culture-based concentrations.) Consistent with procedures for calculating geometric means recommended in the 2012 EPA Recreational Water Quality Criteria (EPA, 2012, p. 39), the geometric mean for a given beach and date was calculated in these cases by first replacing any values outside the range of quantification with the appropriate LOQs. For individual samples involving left-censored data, there is no reliable estimate of the actual concentration but there is strong evidence that it was less than the LLOQ. And for geometric means that were calculated using at least one LLOQ replacement value, there is strong evidence that the geometric mean would have been less than the calculated value if the LLOQs had been low enough so all sample concentrations were quantifiable. Similar reasoning applies to right-censored data, where there is strong evidence that the actual concentration or geometric mean would have been greater than the value involving ULOQ replacements if the ULOQs had been high enough so all sample concentrations were quantifiable. As always in statistical analyses involving censored data, it is important to utilize this “less than” and “greater than” information for censored values. For culture-based data, the only property we use is whether a given estimate or geometric mean is or is not greater than the Michigan standard of 300 *E. coli* per 100 mL. This property is not affected by censoring, because the culture-based LLOQ for Colilert-18 is well below the standard and the ULOQ is well above it. Some of our assessments of the qPCR-based data, however, require that censoring be accounted for.

A related issue is that labs which contributed data were not required to report LOQs for their analyses. This is problematic for qPCR analyses, where LLOQs can vary significantly among sample runs. We were able to obtain all 2019–2020 qPCR LLOQs from the original data workbooks used by the contributing labs, but this was not possible for the 2016–2018 data. Since plausible estimates of LLOQs are essential for some of our assessments of qPCR data, it was necessary to employ substitute LLOQs for the 2016–2018 data. These values were created by random sampling (with replacement) from the upper 50% of the 2019–2020 LLOQs. We originally sampled from all 2019–2020 LLOQs, but visual assessment of the data showed that several of the substitute LLOQs were implausibly low, given their corresponding culture-based concentrations. Restricting sampling to the upper 50% of the 2019–2020 LLOQs partially resolved this problem.

Finally, it is important to be aware of several properties of the data used by Haugland et al. (2021) in deriving the proposed qTV. The data were acquired from samples collected by multiple Michigan labs in 2016–2018 for the state’s beach monitoring program. Most of the samples were split and analyzed by both the collecting lab and EPA’s ORD lab. Samples that were not split (due to cost constraints or technical issues) were analyzed only by the collecting lab. All data generated by Michigan labs that satisfied quality-control criteria were included in the test data set for 2016–2018 used in the present assessment. Most of these data, however, were not used by Haugland et al. (2021) in deriving the proposed qTV: only the EPA data were used for samples that were split and analyzed by both the collecting lab and EPA. Moreover, many samples in the data set used by EPA to derive the proposed qTV were ultimately excluded from their analysis. In particular, samples whose culture-based or qPCR-based concentrations (or both) were censored or classified as statistical outliers were excluded, as were samples from beaches that showed no exceedances of Michigan’s standard of 300 *E. coli* per 100 mL (see Haugland et al., 2021 for further details). The key point is that, while the proposed qTV is based on samples collected for Michigan’s beach monitoring program in 2016–2018, many of the samples were excluded from statistical analyses used in deriving it, and most of the data used were generated by EPA’s ORD lab instead of the collecting labs. Thus, the 2016– 2018 test data utilized in the present assessment are not the same as the 2016–2018 training data utilized in deriving the proposed qTV, though the two sets partially overlap.

### 2.7 Data analysis

As noted in Section 1, the test data employed in our assessment comprise bivariate Colilert-18 and qPCR concentrations from split samples collected and analyzed by multiple labs for Michigan’s annual beach monitoring program during 2016–2018 and 2019–2020. We supplemented these test data with numerical values of three performance measures calculated from results presented by Haugland et al. (2021) for the 2016–2018 training data.

The goals of our assessment are to determine whether (1) qPCR-based *E. coli* concentrations in both sets of test data (2016–2018 and 2019–2020) show good ability to discern the difference between samples that do and do not exceed Michigan’s culture-based standard, (2) the proposed qTV performs similarly and well in predicting culture-based beach notification decisions for both sets of test data, and (3) numerical values of three key performance measures for the proposed qTV on the 2016–2018 and 2019–2020 test data agree closely with the corresponding values on the 2016–2018 training data.

Given our assessment goals, testing statistical null hypotheses of no difference between the two periods of years is not appropriate (similarity does not require equality). Instead, we characterize and compare data and performance measures for the various data sets in multiple ways that allow us to judge whether they are clearly and consistently similar across all measures. If the proposed qTV performs consistently well on the test data, and if numerical values of the key performance measures on the test and training data agree closely, then the proposed qTV is valid in the sense that its performance is good across years and generalizes beyond the data used to derive it.

The main assessment tools we employ are plots of the bivariate data, plots of the empirical distribution functions (EDFs) for actual positive and negative samples, plots of two other empirical functions derived from the EDFs, and a set of numerical performance measures commonly used for assessing decision and classification rules. The numerical performance measures include (among others) the false-positive rate, false-negative rate, and error rate; numerical values of these three measures are reported by Haugland et al. (2021) for the training data or can be calculated from other values they report, and can also be calculated for the test data. Actual positive and negative samples are defined with respect to Michigan’s culture-based standard of 300 *E. coli* per 100 mL: a sample is an actual positive if its culture-based *E. coli* concentration exceeds the standard; otherwise, it is an actual negative. The false-positive rate is the percentage of actual negatives whose qPCR-based concentrations exceed the proposed qTV, the false-negative rate is the percentage of actual positives whose qPCR-based concentrations do not exceed the proposed qTV, and the error rate is the percentage of samples whose implied qPCR-based beach notification decisions do not agree with the culture-based decisions. By EDFs, we mean nondecreasing functions *F*_0_(*x*) and *F*_1_(*x*) that represent the proportions of actual negative and positive samples (respectively) whose log_10_ qPCR-based concentrations are known to be less than or equal to *x* as *x* increases from its minimum to maximum observed values. Plots of *F*_0_(*x*) − *F*_1_(*x*) versus *x*, and of *F*_0_(*x*) versus *F*_1_(*x*) for all *x*, are useful for visually assessing the ability of qPCR-based concentrations to distinguish between actual positive and negative samples (Appendix A, Section A.4). We call *F*_0_(*x*) − *F*_1_(*x*) the discernibility function. Plots of *F*_0_(*x*) versus *F*_1_(*x*) for all *x* are a version of the so-called “receiver operating characteristic” (ROC) curve that is appropriate for data sets that include a non-negligible proportion of left-censored observations of *x*. Standard ROC curves plot *S*_1_(*x*) versus *S*_0_(*x*), where *S*_*i*_(*x*) is the complementary distribution function (survival function) for group *i*, but left-censored data do not permit valid estimates of these functions (briefly, because LLOQs do not provide valid “greater than” information about sample concentrations for left-censored observations but do provide valid “less than” information).

Custom programs written in the R programming language, version 4.2.0 (R Core Team, 2022), were used to create all graphical displays of data and perform all data analyses. Plots of raw data and geometric means for 2016–2018 and 2019–2020 were created with standard R graphics functions. EDFs were computed with R function `ecdf()` and displayed by extracting the relevant components of the object created by this function and then plotting these with standard R graphics functions, thus avoiding the standard plot method for class `ecdf`. The area under each *F*_0_(*x*)-versus-*F*_1_(*x*) curve (AUC) was determined by numerical integration using a custom function written in R.

A set of 18 numerical performance measures were calculated with a custom R program. The measures include, for example, percent true and false positives, percent true and false negatives, percent error rate, true-positive and false-positive rates, true-negative and false-negative rates, and several others (the full list, with synonyms and definitions, is presented with the results in Section 3.2).

## 3 Results

### 3.1 Overall utility of qPCR-based concentrations for identifying actual positive and negative samples

Raw data and geometric means from the 2016–2018 and 2019–2020 test data are plotted in several ways in Fig. 2. Raw data are plotted in the two left columns, geometric means in the two right columns (a guide to interpreting plot types shown in Figs. 2 and 3 is provided in Appendix A, Section A.4).

**Figure 2.**
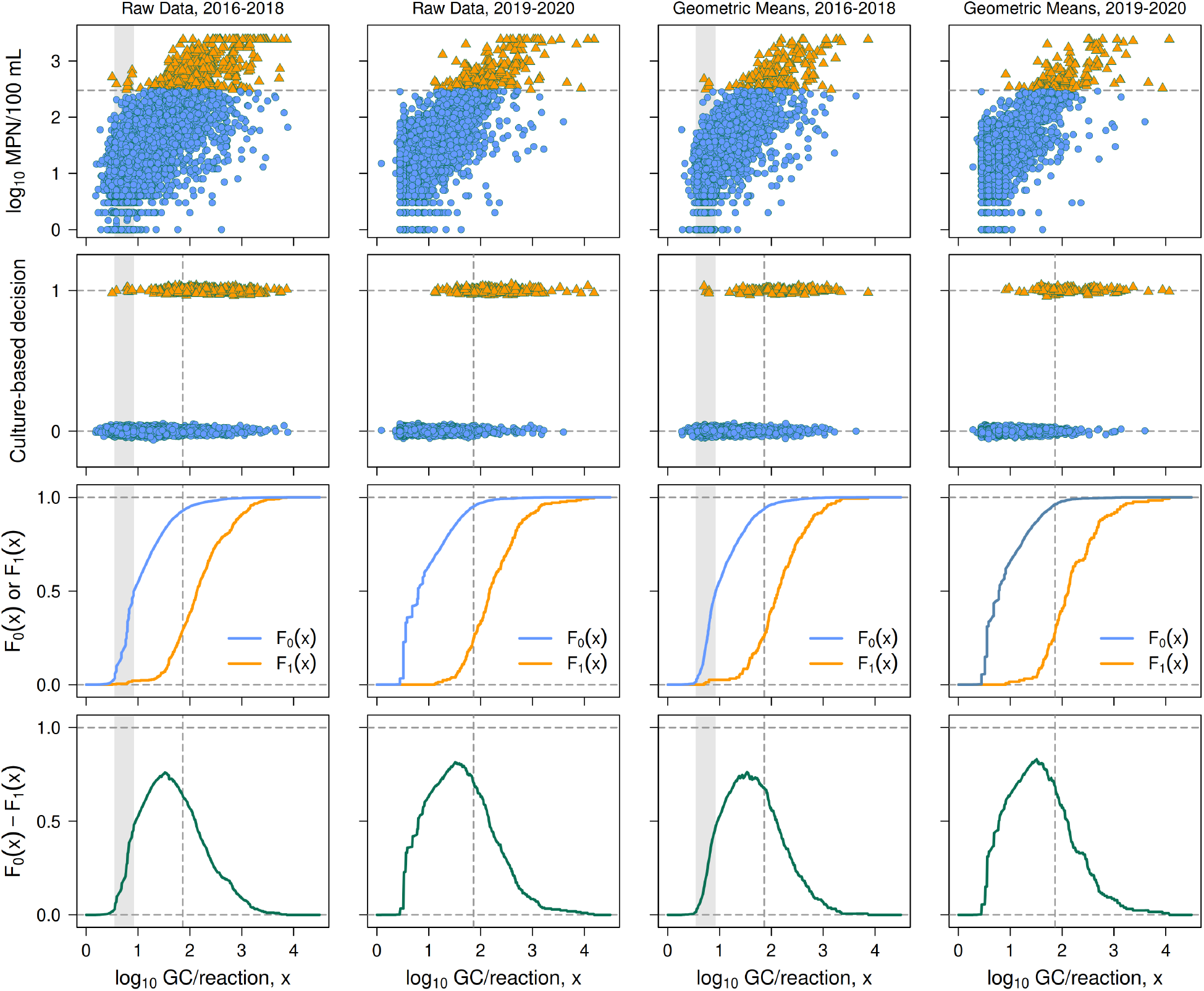
Distributions of actual (culture-based) positive and negative samples for 2016–2018 and 2019–2020 as functions of log_10_ qPCR-based E. coli concentration, compared separately for raw data (two left columns) and geometric means (two right columns). The vertical dashed line in panels of rows 2–4 indicates the proposed Michigan qTV of 1.863 log_10_ GC/reaction. Shaded regions in panels for 2016–2018 data (columns 1, 3) indicate the interval of qPCR-based concentrations containing substitute LLOQs sampled from the known 2019–2020 LLOQs. Row 1: Scatter plots of the paired log_10_-transformed culture-based and qPCR-based concentration data. The horizontal dashed line indicates Michigan’s culture-based beach notification standard of 300 *E. coli*/100 mL (log_10_(300) ≈ 2.48); samples exceeding it are shown as filled triangles, others as filled circles. Row 2: Horizontal strip chart of log_10_ qPCR-based concentrations for culture-based positive and negative samples. Positive samples are plotted at 1 on the vertical axis (with vertical jittering), negative samples at 0. Row 3: Empirical distribution functions *F*_0_(*x*) and *F*_1_(*x*) for actual negative and positive samples. Row 4: Discernibility function *F*_0_(*x*) − *F*_1_(*x*).

**Figure 3.**
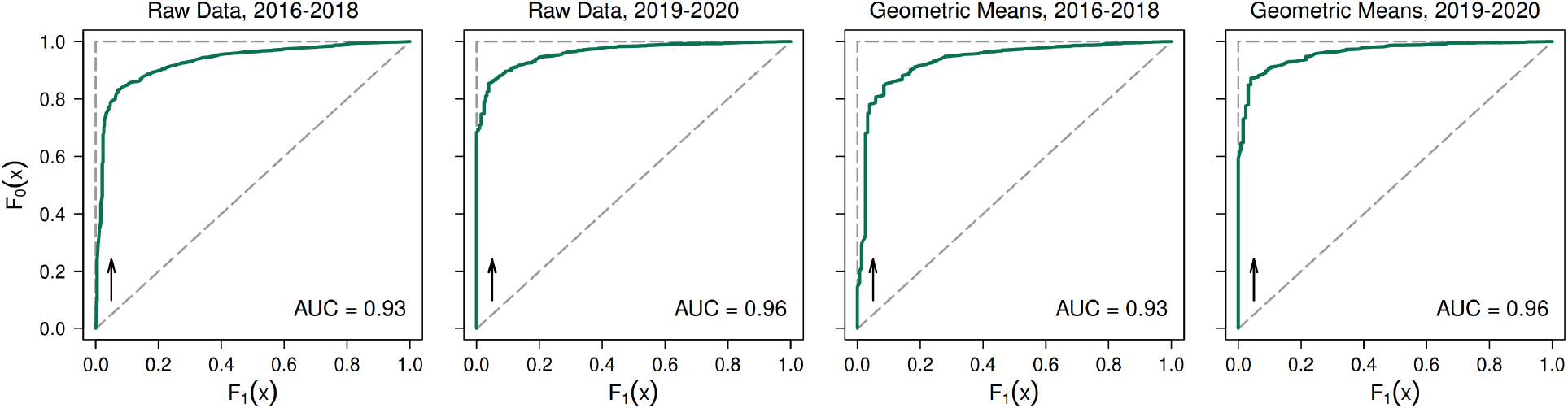
Plots of the EDF for actual negative samples *F*_0_(*x*) versus the EDF for actual positive samples *F*_1_(*x*) over the observed range of log_10_ qPCR-based concentrations *x*. Arrows indicate the direction of increasing *x*. The two left plots are based on raw data, the two right plots on geometric means. The dashed diagonal line indicates the expected relationship if qPCR-based concentrations provide no useful information for distinguishing between actual positive and negative samples. The dashed vertical and horizontal lines together indicate the expected relationship if qPCR concentrations are able to distinguish perfectly between actual positive and negative samples. AUC denotes Area Under the Curve.

Panels in the top row of Fig. 2 show scatter plots of culture-based versus qPCR-based *E. coli* concentrations (both log_10_-transformed) for the various sampling dates in 2016–2018 and 2019–2020. The horizontal dashed line in each panel indicates Michigan’s culture-based standard of 300 *E. coli* per 100 mL (log_10_(300) ≈ 2.48). This decision boundary divides each data set into two subsets: samples that exceed the standard (actual positive samples, denoted by filled triangles) and samples that do not (actual negative samples, denoted by filled circles).

The second row of panels shows these same data in a collapsed form, with actual positive samples (filled triangles) plotted at 1 on the vertical axis and actual negative samples (filled circles) plotted at 0 on the vertical axis. Vertical jittering is used in these strip charts to partially separate the numerous samples with similar or identical qPCR-based concentrations. A small, isolated cluster of points is apparent at the far left of the actual positive samples for 2016–2018. Nearly all of these values involve left-censored data represented by substitute LLOQs. Their separation from the other actual positive samples suggests that the procedure used for choosing substitute LLOQs tended to choose values that were somewhat low, even though sampling was restricted to the upper 50% of the 2019–2020 LLOQs.

If the data did not include censored qPCR-based estimates, it would be appropriate to plot histograms or empirical probability density functions to represent the two groups of samples (actual negative and positive samples) that are evident in the strip charts. But because many of the qPCR-based estimates are left-censored, it is preferable instead to plot EDFs *F*_0_(*x*) and *F*_1_(*x*) for actual negatives and positives. In terms of the strip charts shown in row 2, *F*_0_(*x*) and *F*_1_(*x*) tell us the proportion of actual negatives and actual positives (respectively) lying at or to the left of any given value of *x* on the horizontal axis. The third row of panels in Fig. 2 shows these EDFs for actual negative and positive samples. Note the similar shapes and clear horizontal separation of the two EDFs plotted for each of the four groups of test data, indicating that qPCR-based concentrations allow good (though not perfect) discernment of the difference between actual positive and negative samples for both the 2016–2018 and 2019–2020 test data, regardless of whether one looks at the raw data or the geometric means.

The fourth row of panels in Fig. 2 shows plots of discernibility function *F*_0_(*x*) − *F*_1_(*x*) as a function of log_10_ qPCR-based concentration *x*. The value of this function will be 0 for sufficiently small and large values of *x* and typically will have a single major peak at an intermediate value or range of values. The closer the height of the peak is to 1, the greater is the ability of qPCR-based concentrations to discern the difference between actual positive and negative samples; if the height is identically 1, qPCR-based concentrations permit discernment without error (see the examples in Fig. A.1 of Appendix A). For the four groups of test data, the peak height ranges between roughly 0.76 and 0.83, which are relatively high values indicating good maximum discernibility.

Fig. 3 shows plots of *F*_0_(*x*) versus *F*_1_(*x*) over the entire range of qPCR-based concentrations *x*. If the two distribution functions in row 3 of Fig. 2 showed no separation, we would have *F*_0_(*x*) ≈ *F*_1_(*x*), the graph of *F*_0_(*x*) versus *F*_1_(*x*) would approximate a diagonal line of slope 1, and the area under the *F*_0_(*x*)-versus-*F*_1_(*x*) curve (AUC) would be approximately 0.5 (i.e., half of the plot region, which is a 1-by-1 square). At the opposite extreme, if the two distribution functions in row 3 of Fig. 2 were so well separated that the peak height of the discernibility function was 1, a plot of *F*_0_(*x*) versus *F*_1_(*x*) would approximate a vertical line from the origin to 1 on the *F*_0_ axis (as *F*_0_(*x*) increased to 1 with *F*_1_(*x*) = 0) and then would approximate a horizontal line at *F*_0_ = 1 (as *F*_1_(*x*) increased to 1 with *F*_0_(*x*) = 1), and AUC would be approximately 1. The closer the relationship between *F*_0_(*x*) and *F*_1_(*x*) is to this right-angle curve, and the closer AUC is to 1, the greater is the ability of qPCR-based concentrations to distinguish between culture-based positive and negative samples (see the examples in Fig. A.2 of Appendix A). The panels in Fig. 3 show that the relationship between *F*_0_(*x*) and *F*_1_(*x*) for all four groups of test data is much closer to this right-angle line than to a diagonal line of slope 1, and AUC is greater than 0.9 in all cases. Hosmer et al. (2013, p. 177) present guidelines for interpreting AUC values, according to which values greater than 0.9 indicate “outstanding” ability to discern the difference between actual positive and negative samples. Thus, results for the test data shown in Fig. 3 demonstrate high overall discernibility of qPCR-based concentrations for both the 2016–2018 and 2019–2020 test data, whether one looks at the raw data or the geometric means.

### 3.2 Quantitative performance of Michigan’s proposed qPCR-based criterion

Fig. 4 shows the log_10_-transformed *E. coli* test data for 2016–2018 (left panels) and 2019–2020 (right), plotted with qPCR-based concentration on the horizontal axis and culture-based concentration on the vertical axis. Plots of raw data are shown in the top row and geometric means in the bottom row.

**Figure 4.**
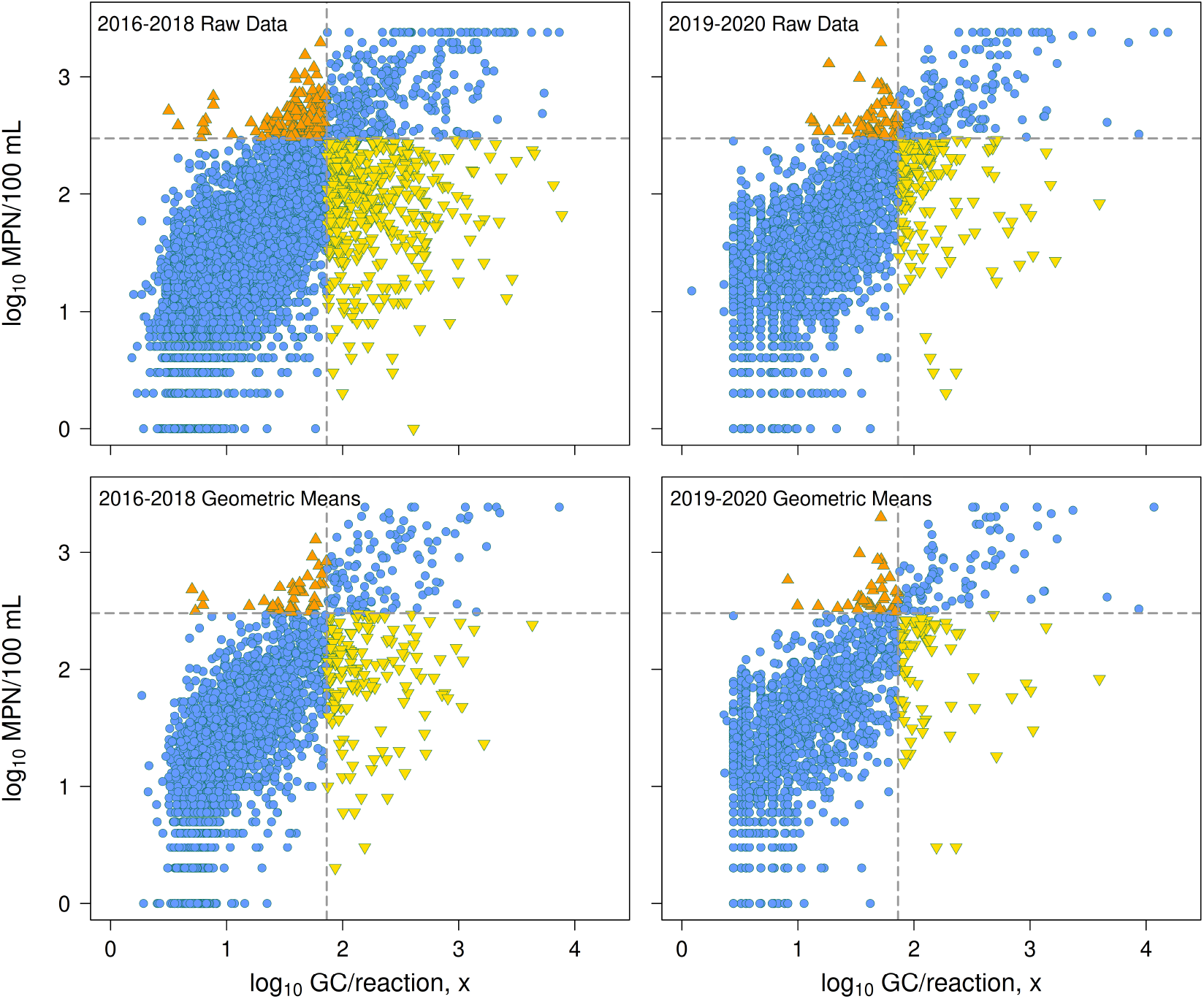
Dual-decision plots of 2016–2018 data (left panels) and 2019–2020 data (right panels), showing both raw data (top panels) and geometric means (bottom panels). Data for 2016–2018 include the same substitute LLOQs as in Figs. 2 and 3. The vertical axis in each panel represents culture-based concentration while the horizontal axis represents qPCR-based concentration, both on a log_10_ scale. The horizontal dashed line in each panel indicates Michigan’s culture-based standard of 300 *E. coli*/100 mL (log_10_(300) ≈ 2.48); the vertical dashed line in each panel indicates Michigan’s proposed qTV 1.863 log_10_ GC/reaction. Filled circles: samples where qPCR-based decisions agree with culture-based decisions (true-positive and true-negative samples). Upright triangles: false-negative samples. Inverted triangles: false-positive samples.

Each panel in Fig. 4 has a horizontal dashed line at Michigan’s culture-based standard of 300 *E. coli* per 100 mL (log_10_(300) ≈ 2.48) and a vertical dashed line at Michigan’s proposed qTV of 1.863 log_10_ GC/reaction. Michigan’s recreational WQS specify that a beach notification should be issued when the reported culture-based *E. coli* concentration for a particular beach on a particular day exceeds 300 *E. coli* per 100 mL. If the proposed qTV is used instead, a beach notification should be issued when the reported log_10_-transformed qPCR-based concentration exceeds 1.863 log_10_ GC/reaction. Thus, the dashed lines in each panel represent two different decision boundaries.

These boundaries divide each panel into four quadrants (for assessment purposes, we apply the boundaries to both raw data and geometric means). In the lower left and upper right quadrants, qPCR-based decisions agree with culture-based decisions. Viewing the culture-based decisions as correct (in the sense that they are the decisions mandated by state law), the qPCR-based decisions in the lower left quadrant are true negatives while those in the upper right quadrant are true positives (all shown as filled circles). Data in the lower right quadrant are false positives (inverted triangles), where the implied qPCR-based decision is to issue a beach notification but the correct, culture-based decision is not to issue one. Data in the upper left quadrant are false negatives (upright triangles), where the implied qPCR-based decision is not to issue a beach notification but the correct, culture-based decision is to issue one; these cases represent the most serious type of decision error from a human health perspective. This figure visually demonstrates the overall similarity of the distributions of data among the four quadrants for the four groups of test data. Numerical percentages of samples in the four quadrants are also very similar across the four groups (Table 1, rows 1–4).

**Table 1.**
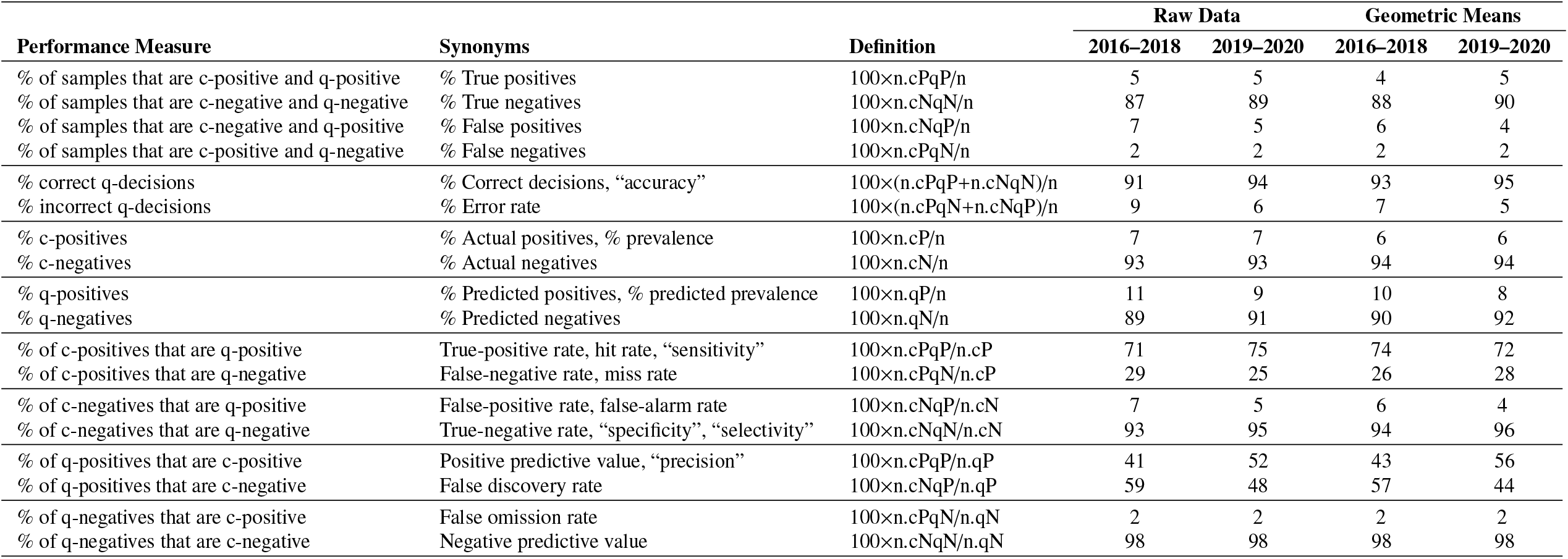
Comparison of quantitative performance measures for Michigan’s proposed qPCR threshold value. Explicit names for the performance measures are stated in column 1, common synonyms in column 2, and definitions in column 3. Synonyms in quotation marks are misleading and should be avoided (see Appendix A, Section A.5). Performance measures are divided by horizontal lines into groups of exhaustive and mutually exclusive classes whose percentages sum to 100 (when not rounded to integers). c-decision: a decision based on a culture-based concentration; q-decision: a decision based on a qPCR-based concentration; c-positive: culture-based concentration > 300 *E. coli*/100 mL; c-negative: culture-based concentration ⩽ 300 *E. coli*/100 mL; q-positive: qPCR-based concentration > 1.863 log_10_ GC/reaction; q-negative: qPCR-based concentration ⩽ 1.863 log_10_ GC/reaction; n.cPqP: number of samples that are c-positive and q-positive; n.cPqN = number of samples that are c-positive and q-negative; etc.; n.cP = n.cPqP + n.cPqN = total number of c-positive samples; etc.; n = total number of samples.

Numerical values of several measures for assessing the performance of Michigan’s proposed qTV and for comparing this performance in the two data sets are presented in Table 1. The basic ideas behind these measures are simple and intuitive: good performance of the proposed qTV means that beach notification decisions based on it agree with the culture-based decisions in a high percentage of cases and disagree in a low percentage of cases. In using these numerical measures to evaluate the performance of the proposed qTV, it is important to remember that qPCR-based beach notification decisions depend only on whether the reported concentration estimates exceed the proposed threshold of 1.863 log_10_ GC/reaction. Because all the known LLOQs from 2019–2020 are well below this threshold, and because it is highly likely that all the unknown LLOQs in 2016–2018 also were well below it, neither the prevalence of left-censored qPCR-based concentrations nor our procedure for choosing substitute LLOQs affects the numerical performance measures we report.

Most of the measures listed in Table 1 are commonly used in assessing the performance of decision or classification rules in various applications, especially medical diagnostic tests (e.g., Webb and Copsey, 2011; Larner, 2021). In several cases, the same measure is referred to by different names in different applications. More importantly, some of the common synonyms actually do not represent the property that the name implies and are therefore misleading. We address the issues of duplicate and misleading terminology by identifying (in column 1 of the table) each performance measure by an explicit name that is directly related to the definition of the measure (stated in column 3), then listing (in column 2) common synonyms we are aware of, with misleading synonyms enclosed in quotation marks. For reasons discussed in Section A.5 of Appendix A, we strongly recommend that use of these misleading terms be avoided; they are included in Table 1 for the benefit of readers who may be familiar with the respective performance measures in other applications under these unfortunate names.

Numerical values of the performance measures in Table 1 indicate that the performance of Michigan’s proposed qTV is highly consistent between the 2016–2018 and 2019–2020 test data sets for both the raw data and the geometric means. This result is important, because all data used in deriving the proposed qTV of 1.863 log_10_ GC/reaction came from the 2016–2018 period. The high level of consistency also applies to the test data versus the training data. Table 2 displays the false-positive, false-negative, and error rates for the training data from 2016–2018, along with the corresponding rates for four groups of test data. All values for the test data closely resemble those for the training data and in fact suggest a tendency toward slightly lower error rates in the test data. The consistency of performance between time periods and between training and test data is evidence for the validity of the proposed qTV, since it shows that the threshold performs similarly on data from the period for which it was developed and on data from a different period, and on both training and test data.

**Table 2.** Comparison of false-positive rates (FPR), false-negative rates (FNR), and error rates (ER) for training data and four groups of test data. Performance measures are defined in Table 1.

| Data | FPR (%) | FNR (%) | ER (%) | Source |
| --- | --- | --- | --- | --- |
| Training data: |  |  |  | Haugland et al. (2021) |
| 2016–2018 (39 sites) | 7 | 30 | 10 |  |
| Test data: |  |  |  | This study |
| 2016–2018 raw data | 7 | 29 | 9 |  |
| 2016–2018 geometric means | 6 | 26 | 7 |  |
| 2019–2020 raw data | 5 | 25 | 6 |  |
| 2019–2020 geometric means | 4 | 28 | 5 |  |

Values of the performance measures in Table 1 also indicate that performance of the proposed qTV is not only consistent between data sets but is consistently good. In particular, we note that qPCR-based decisions are correct in 91–95% of samples (Table 1: % correct decisions), the error rate is 5–9%, negative qPCR-based decisions are consistently correct in 98% of cases (negative predictive value), the true-positive rate is 71–75%, and the true-negative rate 93–96%. The false-negative rate is 25–29%, which is higher than the false-positive rate (4–7%) but applies to a much smaller percentage of samples (6–7% versus 93–94%). As noted above, Haugland et al. (2021) found similar false-negative, false-positive, and error rates for the training data. In studies at other geographic locations, Kephart and Bushon (2009) obtained false-negative rates of 9–100%, false-positive rates of 0–21%, and error rates of 6–25% for paired qPCR-based and culture-based *E. coli* concentrations in split samples from two Lake Erie beaches in Cleveland, Ohio, while Shrestha and Dorevitch (2019) obtained false-negative and false-positive rates of 18% and 20% (respectively) for paired analyses in split samples from eight Lake Michigan beaches in Chicago, Illinois (both studies derived their own region-specific qTVs).

## 4 Discussion

The goals of this study are to determine whether (1) qPCR-based *E. coli* concentrations consistently show good ability to distinguish between actual positive and negative samples in test data sets from 2016–2018 and 2019–2020, (2) Michigan’s proposed qTV performs consistently well in predicting culture-based beach notification decisions for both sets of test data, and (3) the proposed qTV performs consistently well on the test and training data, as judged by three numerical performance measures that could be calculated for both types of data.

With regard to the first goal, the results of our assessment show that the ability qPCR-based *E. coli* concentrations to distinguish between actual positive and negative samples in the test data is very good (“outstanding”, on the scale of Hosmer et al., 2013). This conclusion holds for both the 2016–2018 and 2019–2020 test data, regardless of whether one looks at the raw data or geometric means.

Regarding the second goal, the results show that Michigan’s proposed qTV performed well, with an error rate consistently less than 10%, a true-negative rate consistently greater than 90% (meaning that over 90% of actual negatives were correctly identified as negatives by qPCR), and a negative predictive value that was consistently 98% on the test data (meaning that 98% of samples identified as negatives by qPCR were actual negatives; equivalently, only 2% were actual positives). This conclusion again holds for both the 2016–2018 and 2019–2020 test data, whether one looks at the raw data or geometric means.

Regarding the third goal, the results show that the false-positive, false-negative, and error rates are very similar for the training data from 2016–2018, the raw data and geometric means for test data from 2016–2018, and the the raw data and geometric means for test data from 2019–2020. The false-positive rates ranged from 4–7%, the false-negative rates from 25–30%, and the error rates from 5–10%, with the training data exhibiting the highest value of each measure of decision error and the test data exhibiting all the lowest values. Thus, the proposed qTV performed at least as well on the test data as on the training data.

Taken together, the results of our assessment indicate that qPCR-based *E. coli* concentrations show good ability to distinguish between actual positive and negative samples, that the proposed qTV performed consistently well when thoroughly evaluated on the 2016–2018 and 2019–2020 test data, and that it also performed at least as well on the test data as on the training data when assessed using three measures of decision error whose values could be calculated for all data sets. These results support the proposed qTV as a valid alternative to Michigan’s culture-based standard.

It is perhaps not surprising that the proposed threshold performed well on the 2016–2018 test data, since it was developed using data from this period (though not exactly the same data). However, none of the samples or data for 2019–2020 were used to develop the threshold. Moreover, most of the training data for 2016–2018 were produced by a single lab (EPA’s ORD lab), whereas the 2016–2018 and 2019–2020 test data sets were produced by multiple labs across the state of Michigan. For these reasons, the fact that the proposed qTV performed similarly on the 2016–2018 training data and the 2016–2018 and 2019–2020 test data is compelling evidence for the validity of both the proposed threshold and the procedure of using multiple labs to collect samples and perform the qPCR analysis.

An interesting feature of the discernibility functions in Fig. 2 is that maximum discernibility consistently occurs at qPCR-based concentrations that are somewhat lower than Michigan’s proposed qTV of 1.863 log_10_ GC/reaction. This result mainly reflects the fact that there is more than one plausible approach to choosing an optimal decision rule for beach notifications, and each approach yields a somewhat different decision boundary. The proposed Michigan criterion is based on a statistical calibration approach that seeks to characterize the relationship between non-censored qPCR-based and culture-based *E. coli* concentrations (Haugland et al., 2021) instead of maximizing discernibility of actual positive and negative samples as defined here. The calibration approach requires that both the culture-based and qPCR-based concentrations for every sample be represented by measured values, implying that all samples for which either concentration was censored must be excluded. This exclusion removes a large number of censored observations (roughly half of all samples, mostly left-censored) that provide valid “less than” or “greater than” concentration information that is useful in discerning the difference between actual positive and negative samples.

Some of the effects of excluding censored samples on the utility of qPCR-based concentrations for identifying actual positive and negative samples are illustrated in Fig. 5. It shows plots of the EDFs, discernibility functions, and *F*_0_(*x*)-versus-*F*_1_(*x*) relationships for the 2019–2020 data with censored observations excluded (solid lines) and included (dotted lines). In the latter case, censored observations are represented by their LLOQs, which are known for the 2019–2020 data. Note that excluding censored observations shifts the EDF for actual negative samples substantially to the right but has little effect on the EDF for actual positive samples. It also shifts the discernibility peak to the right and reduces its height, lowers the *F*_0_(*x*)-versus-*F*_1_(*x*) curve, and reduces AUC from 0.96 to 0.91. The effect of excluding censored observations, then, is to reduce the ability of qPCR-based concentrations to discern the difference between actual positive and negative samples, and also to move the qPCR-based concentration at which maximum discernibility occurs to the right, closer to the proposed qPCR-based criterion (vertical dashed line) but still not coinciding with it. Thus, part of the difference between the location of the discernibility peak and the proposed qPCR-based criterion can be attributed to the fact that the calibration approach discards information contained in censored observations, while the remainder of the difference can be attributed to the use of different optimality criteria.

**Figure 5.**
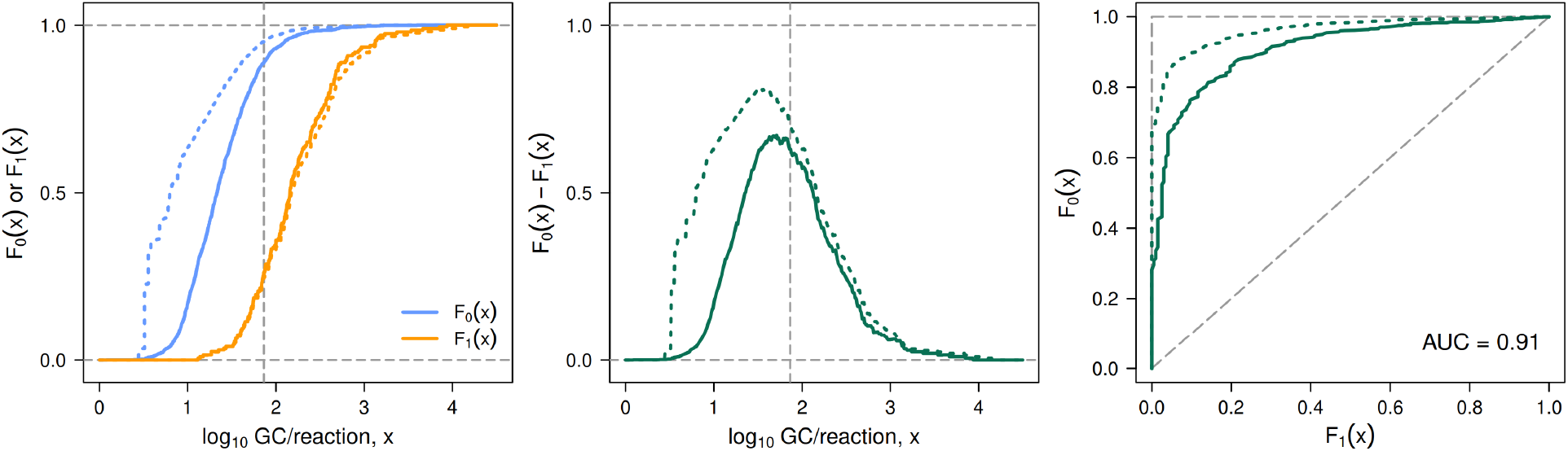
Effects of excluding censored samples on the utility of qPCR-based *E. coli* concentrations for distinguishing between actual positive and negative samples, based on raw test data for 2019–2020. Left: Empirical distribution functions *F*_0_(*x*) for actual negative samples and *F*_1_(*x*) for actual positive samples as functions of log_10_ qPCR-based concentration *x* when samples with censored concentrations are excluded (solid lines) or included (dotted lines). Center: Discernibility *F*_0_(*x*) − *F*_1_(*x*) as a function of log_10_ qPCR-based concentration *x* when censored samples are excluded (solid line) or included (dotted line). Right: *F*_0_(*x*) versus *F*_1_(*x*) for all qPCR-based concentrations *x* when censored samples are excluded (solid line) or included (dotted line). AUC is the area under the curve. For comparison, AUC = 0.96 when censored samples are included (see Fig. 3).

Table 1 shows that 89–92% of samples from Michigan beaches were qPCR-based negatives, of which 98% were confirmed to be actual negatives. These results indicate that negative qPCR-based decisions are highly but not perfectly reliable indications that the culture-based decision would also be negative if both types of analysis were performed. But as pointed out in Section 3.2, 25–29% of the actual positives were qPCR-based negatives. Thus, although the proposed qPCR-based beach notification criterion performs very well in correctly detecting actual negatives, its performance is not as impressive in correctly detecting actual positives. The relatively high false-negative rate is not as serious a problem as it might at first appear, because actual positives represent only 6–7% of samples and thus are relatively rare. It must also be remembered that the fundamental reason for developing a qTV is that culture-based concentration estimates apply to yesterday’s conditions while qPCR-based estimates apply to today’s. Given that concentrations of fecal indicator bacteria at a beach can change markedly from one day to the next (Whitman et al., 1999; Boehm et al., 2002; Whitman and Nevers, 2004; Boehm, 2007), a relatively high false-negative rate for rarely-occurring positive samples may be viewed as an acceptable trade-off for gaining the advantage of same-day concentration estimates.

The rarity of actual positives at bathing beaches and the fact that false-negative errors were not treated as being more serious than false-positive errors in deriving Michigan’s proposed qTV are two of the main reasons that the proposed threshold yields a higher false-negative rate (25–29%) than false-positive rate (4–7%). If false-negative errors were weighted more heavily than false-positive errors to reflect their greater importance for human health, one would expect the optimal qPCR-based decision boundary to decrease and thus produce fewer false negatives. But as can be seen in any of the panels in the second or third row of Fig. 2, decreasing the decision boundary also increases the number of false positives and hence the number of unnecessary beach advisories or closings.

Another contributor to the relatively high false-negative rate is that certain beaches repeatedly show false negatives. Thus, there seems to be a currently unknown source of variability among beaches that affects this type of decision error, with certain beaches being particularly prone to it. If the source of this variability can be identified, it might be possible to measure it and use its value in some way (e.g., to stratify beaches or as a statistical covariate) to decrease the false-negative rate without increasing the false-positive rate. More work clearly needs to be done on this issue. In this regard, we note that the methods of data analysis and visualization we have employed are useful for identifying individual beaches that repeatedly yield erroneous qPCR-based decisions in regional and local data sets. Once these exceptional beaches have been identified, they can be targeted for follow-up assessments to help understand the causes of the erroneous decisions (e.g., qPCR inhibition, specific landscape properties, temporal variables such as rain events and algal blooms). Nevertheless, results of the visual and quantitative assessments we have presented show that Michigan’s proposed qTV for beach notifications based simply on qPCR-based *E. coli* concentrations performs well and, in particular, performs similarly well for the 2016–2018 training data and both the 2016–2018 and 2019–2020 test data.

## Acknowledgements

We thank Rich Haugland from EPA’s Office of Research and Development (Cincinnati, Ohio) for his encouragement and insightful comments, the many students and technicians who helped produce the data underlying this assessment, and the Michigan Network for Environmental Health and Technology

## A Appendix

### A.1 Relevant Michigan water-quality standards for total-body-contact recreation

Monitoring of beaches in Michigan is voluntary and is conducted by local health departments, which are required to comply with state water-quality standards (WQS) for total-body-contact recreation according to R 333.12544 of the Public Health Code, 1978 PA 368 (Act 368), as amended. The WQS require that “at no time shall the waters of the state protected for total-body-contact recreation contain more than a maximum of 300 *E. coli* per 100 mL. Compliance shall be based on the geometric mean of 3 or more samples taken during the same sampling event at representative locations within a defined sampling area” (R 323.1062 Microorganisms, Rule 62(1), of the Part 4 rules, WQS, promulgated under Part 31, Water Resources Protection, of the Natural Resources and Environmental Protection Act, 1994 PA 451, as amended).

### A.2 List of contributing laboratories

**Table A.1.**
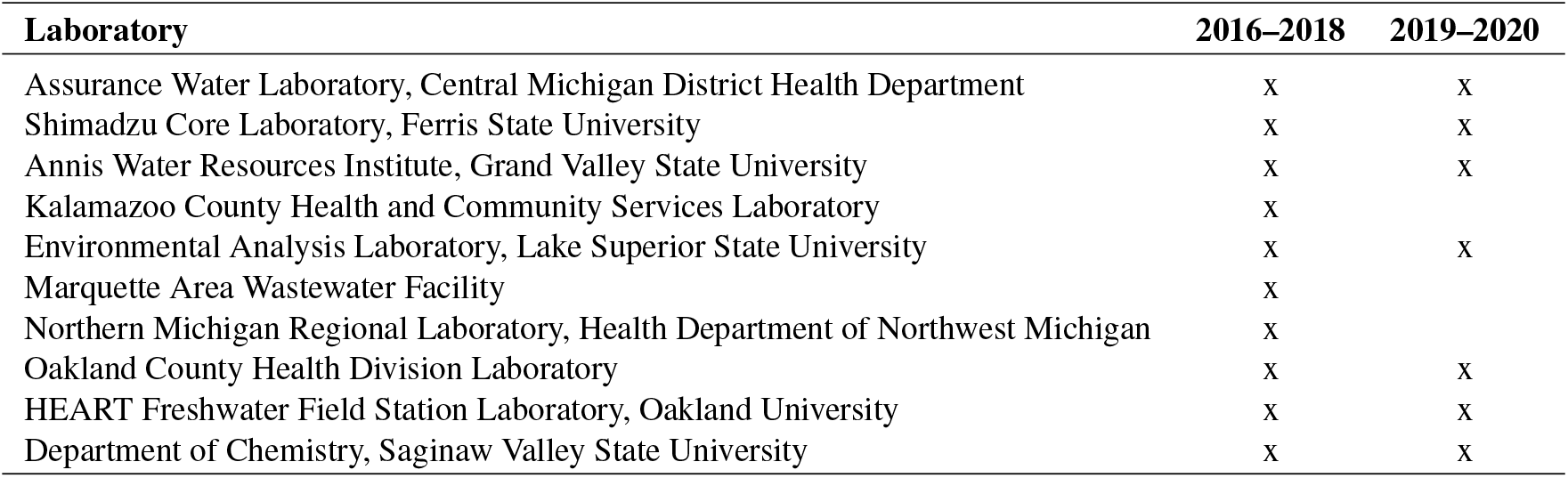
Laboratories which contributed Colilert-18^®^ and qPCR data to the 2016–2018 or 2019–2020 Michigan beach monitoring programs that were used in the present assessment of the proposed qPCR threshold value. An “x” indicates that the respective laboratory contributed data.

| Laboratory | 2016–2018 | 2019–2020 |
| --- | --- | --- |
| Assurance Water Laboratory, Central Michigan District Health Department | x | x |
| Shimadzu Core Laboratory, Ferris State University | x | x |
| Annis Water Resources Institute, Grand Valley State University | x | x |
| Kalamazoo County Health and Community Services Laboratory | x |  |
| Environmental Analysis Laboratory, Lake Superior State University | x | x |
| Marquette Area Wastewater Facility | x |  |
| Northern Michigan Regional Laboratory, Health Department of Northwest Michigan | x |  |
| Oakland County Health Division Laboratory | x | x |
| HEART Freshwater Field Station Laboratory, Oakland University | x | x |
| Department of Chemistry, Saginaw Valley State University | x | x |

### A.3 qPCR methods

#### A.3.1 qPCR analysis

Methods of qPCR analysis followed EPA Draft Method C (Aw et al., 2019; Lane et al., 2020; Sivaganesan et al., 2019). Briefly, two 100-mL aliquots from each individual or composite sample were filtered through a 0.4-*µ*m pore size polycarbonate filter (Millipore or equivalent), after which 20 mL of Phosphate Buffer Saline (PBS) (pH = 7.4 ± 0.2) was filtered as a rinse step and stabilizer. The filter was then collected using aseptic techniques and placed in a 2.0-mL microcentrifuge tube containing 0.3 ± 0.01 g of acid-washed glass beads (212-300 *µ*m, Sigma G1277-500G). In 2016–2018, one filter was processed by the collecting lab and the other was sent to EPA’s Office of Research and Development (ORD) lab (Cincinnati, Ohio) for analysis; samples from 2019–2020 were analyzed by the collecting lab only. Samples were then either processed immediately or stored at 80 °C until qPCR analysis (short-term storage at −20 °C was permitted for labs unable to store at −80 °C). As positive and negative controls, a calibrator filter was prepared using 1.0 mL of a 1 × 10^4^ cells/mL suspension of *E. coli* from MultiShot-10E8 BioBalls™ (BioMèrieux, Lombard, Illinois, Reference #56146) in sterile PBS, and a filter blank was prepared by filtering 20 mL PBS.

DNA extraction consisted of a crude lysis method described by Aw et al. (2019). Briefly, 600 *µ*L of Qiagen AE buffer containing 0.2 *µ*g/mL of salmon DNA (SAE) was added to each tube to resuspend the filter, after which the tubes were homogenized at 5 m/s or bead milled at 5000 reciprocations/min for 1 min. Samples were then centrifuged for 1 min at 12,000 × *g* to pellet any debris, 400 *µ*L of the supernatant was transferred to a clean microcentrifuge tube and centrifuged for an additional 5 min, and 350 *µ*L of the supernatant was transferred to another clean microcentrifuge tube for qPCR analysis.

qPCR analysis was performed using TaqMan™ hydrolysis probes (Thermo Fisher Scientific, Life Sciences Group, Carlsbad, CA). The qPCR assays used for analysis were *E. coli* (EC23S857) and salmon DNA (Sketa22) (Chern et al., 2011; EPA, 2012); sequences are provided in Table A.2. Salmon DNA served as a sample processing control (SPC), and each qPCR assay was run as a singleplex. Three controls (positive, negative, and no-treatment control consisting of Type 1 water) were analyzed on each plate. Assay mixes were prepared using confirmed *E. coli*-free TaqMan Environmental MM 2.0 (Thermo Fisher Scientific, Grand Island, New York) (12.5 *µ*L), 2.5 *µ*L of 2.0 mg/mL bovine serum albumin (BSA) from fraction V powder (Sigma B-4287 or equivalent), 3.0 *µ*L of primer-probe mix (a concentration of 1 *µ*M of each primer) and 80 nM of probe in the reactions, 2.0 *µ*L of DNA-pure water, and 5.0 *µ*L of extracts, for a final reaction volume of 25 *µ*L. Samples and controls were plated in singleplex and analyzed on Applied Biosystems StepOnePlus™ thermal cyclers; thermal cycler conditions are provided in Section A.3.3. The detection limit was manually set to 0.03 ΔRn.

Results from qPCR analysis were quantified for *E. coli* using a 5- or 6-point standard curve from plasmid-derived samples. Standards were prepared and quantified by EPA’s ORD lab in Cincinnati, Ohio, as described by Sivaganesan et al. (2019). Standard curves and concentration estimates were calculated automatically by Draft Method C Excel^®^ workbooks. Each lab entered its data in these workbooks for each sample season. The workbooks screened for quality control and, if all conditions were satisfied, calculated *E. coli* concentrations as log_10_ gene copies per reaction, using the standard curve with adjustments for DNA recovery provided by the SPC assay results (Lane et al., 2020). LLOQs were determined separately for each standard curve (Lane et al., 2020).

#### A.3.2 Sequences

**Table A.2.**
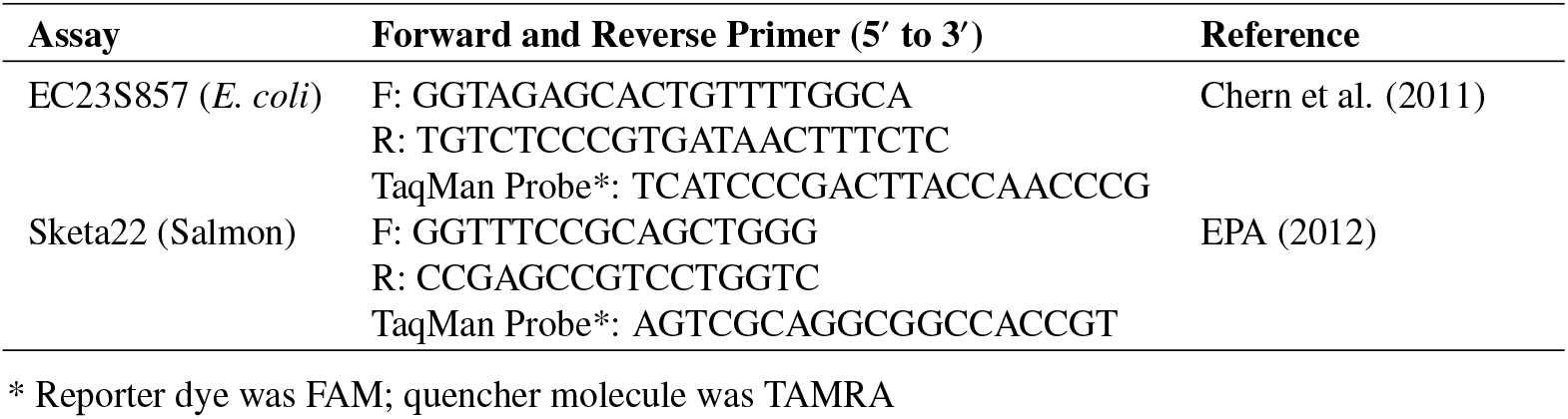
Draft Method C *E. coli* and salmon DNA primer sequences and TaqMan™ probe sequences.

#### A.3.3 Thermal cycler conditions

Thermal cycler parameters included an initial holding stage (50.0 °C, 2 min; 95.0 °C, 10 min), followed by 40 cycles of DNA denaturation and primer/probe annealing (95.0 °C, 15 s; 56.0 °C, 60 s). Fluorescence was measured (StepOnePlus™, Applied Biosystems) at the end of each cycle. The fluorescence threshold was manually set to 0.03 ΔRn, and baseline cycles were set to AUTO determination (Sivaganesan et al., 2019).

### A.4 Examples illustrating the interpretation of some useful diagnostic plots

Here we provide examples based on simulated data to illustrate the interpretation of various diagnostic plots that are useful for assessing the ability of qPCR-based concentrations to distinguish between actual positive and negative samples. The examples consist of three simulated data sets and corresponding visual diagnostic plots. In the first data set, there is no correlation between qPCR-based and culture-based concentrations, and qPCR-based concentrations are therefore useless for distinguishing between actual positive and negative samples. In the second, there is strong positive correlation between the two types of concentrations, and actual positive and negative samples can be distinguished reasonably well on the basis of qPCR-based concentrations but not without error. In the third example, qPCR-based concentrations are able to distinguish between actual positive and negative samples without error. Simulated data were generated with function `rmvnorm()` from R package mvtnorm version 1.1-1 (Genz and Bretz, 2009; Genz et al., 2020). No censoring was applied to the data. All analyses and plots of data were performed using the R programming language and computation environment, version 4.2.0 (R Core Team, 2022).

Fig. A.1 shows the three simulated data sets and various diagnostic plots created from them. Each column of panels in the figure corresponds to one of the three simulated data sets. The first column corresponds to data where there is no correlation between qPCR-based and culture-based concentrations, the second to data where there is strong positive correlation that permits reasonably good discernment of the difference between actual positive and negative samples, and the third to data where actual positive and negative samples can be distinguished without error on the basis of qPCR-based concentration.

Panels in the first row of Fig. A.1 show the three data sets. The second row shows the same data but with a horizontal dashed line, analogous to Michigan’s culture-based recreational WQS for *E. coli*, that divides the data into actual positive and negative samples (red and blue dots, respectively). The third row shows horizontal strip charts for each data set, with actual positive samples (red dots) plotted at 1 on the vertical axis (with vertical jittering) and actual negative samples (blue dots) plotted at 0. Note that these strip charts give a good visual sense of the degree to which the actual positive and negative samples are separated horizontally along qPCR-based concentration axis, which in turn is a good indicator of the ability of qPCR-based concentrations to discern the difference between actual positive and negative samples.

Plots in the first three rows of Fig. A.1 are simply alternative ways of displaying the data. Plots in the four subsequent rows show different functions, defined on the data, that visually reveal the ability of qPCR-based concentrations to distinguish between actual positive and negative samples.

The fourth row shows empirical probability density functions *f*_1_(*x*) and *f*_0_(*x*) for actual positive and negative samples, created using standard R function `density()`. Like the horizontal strip charts, these functions are useful for giving a visual sense of the degree of horizontal separation between actual positive and negative samples along the qPCR-based concentration axis. Estimation of empirical density functions, however, is problematic for data sets that include a substantial proportion of censored observations, as is true of the Michigan beach-monitoring data.

The fifth row shows empirical (probability) distribution functions (EDFs) *F*_1_(*x*) and *F*_0_(*x*) for actual positive and negative samples, created using standard R function `ecdf()`. Here, *F*_*i*_(*x*) represents the proportion of samples in group *i* (0: actual negative samples, 1: actual positive samples) whose log_10_ qPCR-based concentrations are less than or equal to *x*. (Especially in applied statistics, probability distribution functions are sometimes called cumulative probability distribution functions. But as probability theorists often point out, “cumulative” is redundant because probability distribution functions are cumulative by definition.) In terms of the strip charts shown in row 3, *F*_0_(*x*) and *F*_1_(*x*) represent the proportion of the blue dots and red dots (respectively) lying at or to the left of any given value of *x* on the horizontal axis. Each of these func-tions has a vertical jump at each distinct qPCR-based concentration, including left-censored values. Unlike empirical density functions, EDFs are appropriate for data sets with a substantial proportion of left-censored observations, because we know that the actual concentration for a left-censored observation is less than or equal to the LLOQ.

**Figure A.1.**
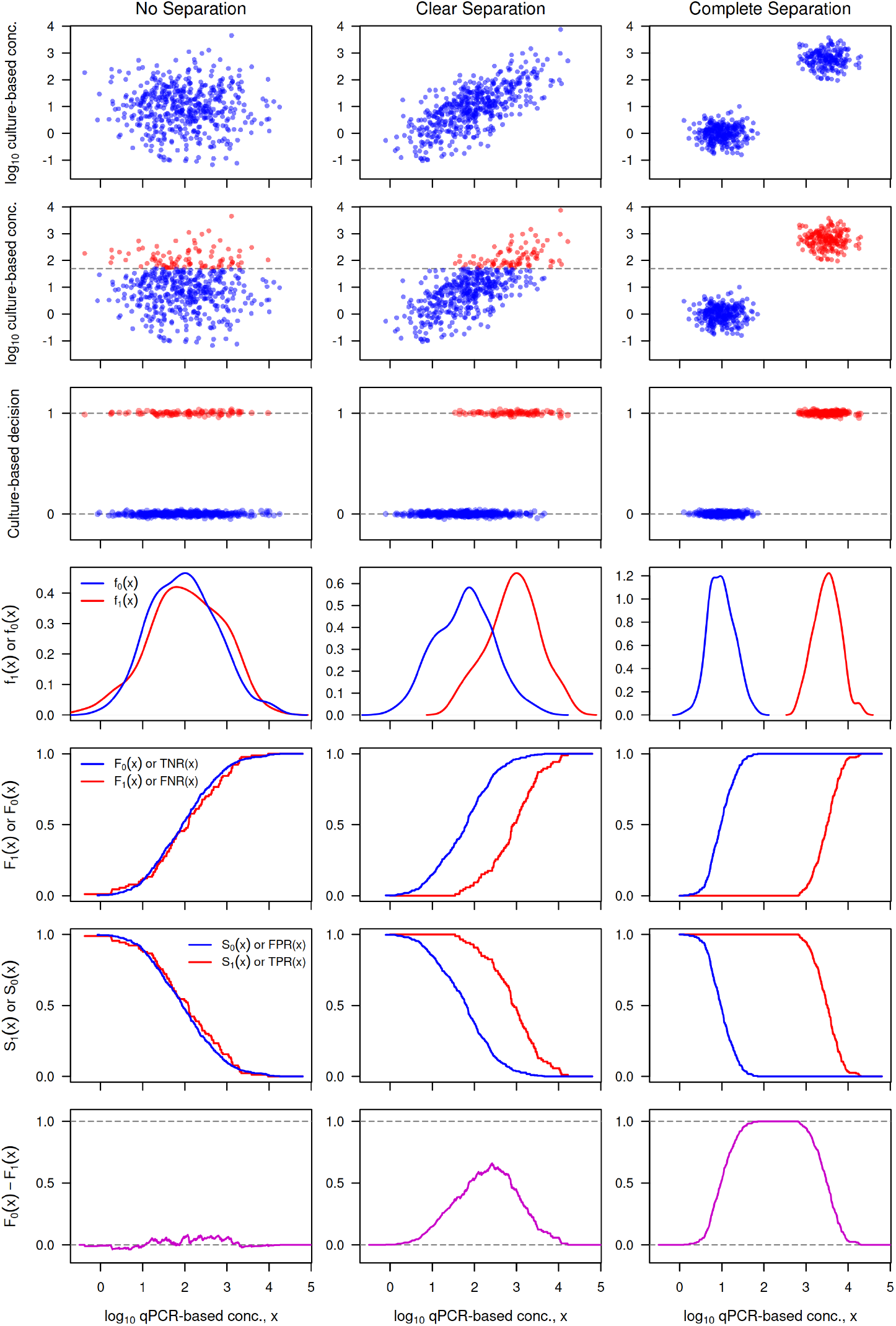
Plots of three simulated data sets (top row) and various visual diagnostics for assessing the utility of qPCR-based concentrations for distinguishing between actual positive and negative samples (rows 2–7). Each column corresponds to one data set. The dashed horizontal line in panels of row 2 is analogous to Michigan’s culture-based recreational WQS for *E. coli*. Notation: *f*_1_(*x*) and *f*_0_(*x*) denote empirical probability density functions for actual positive and negative samples, *F*_1_(*x*) and *F*_0_(*x*) denote the corresponding empirical distribution functions, *S*_1_(*x*) and *S*_0_(*x*) denote the corresponding empirical complementary distribution functions (survival functions), and *F*_0_(*x*) − *F*_1_(*x*) denotes the discernibility function.

The sixth row shows the corresponding empirical survival functions (complementary distribution functions) *S*_1_(*x*) and *S*_0_(*x*), where *S*_*i*_(*x*) = 1 − *F*_*i*_(*x*) represents the proportion of samples in group *i* whose log_10_ qPCR-based concentrations are greater than *x*. These functions are not appropriate for data sets with a substantial proportion of left-censored observations, because their “greater than” property is not consistent with the “less than” information which left-censored observations provide.

Plots in the final row show discernibility function *F*_0_(*x*) − *F*_1_(*x*). The difference *F*_0_(*x*) − *F*_1_(*x*) as a function of *x* is a useful measure of the degree to which qPCR-based concentrations permit discernment of the difference between actual positive and negative samples. If the strips of actual positive and negative samples in row 3 of Fig. A.1 showed no overlap (as is true in column 3), the qPCR-based concentration for any given sample would tell us without error which group a sample belongs to. For qPCR concentrations below the lowest observed concentration, we would have *F*_0_(*x*) = 0 and *F*_1_(*x*) = 0 and therefore *F*_0_(*x*) − *F*_1_(*x*) = 0. At somewhat higher qPCR concentrations, the EDF *F*_0_(*x*) for negative samples would increase step-wise to 1 while the corresponding function *F*_1_(*x*) for positive samples remained at 0 (because no positive samples would have such low qPCR-based concentrations). Thus, the difference *F*_0_(*x*) − *F*_1_(*x*) would increase from 0 to 1, its maximum possible value. With further increases in *x*, the value of *F*_0_(*x*) would remain at 1, qPCR-based concentrations for positive samples would eventually be encountered, *F*_1_(*x*) would then increase step-wise to 1, and *F*_0_(*x*) − *F*_1_(*x*) would decrease back to 0. Thus, the difference *F*_0_(*x*) − *F*_1_(*x*) would be 0 at sufficiently small and large values of *x* but would be 1 for some intermediate value or range of values. (Note that all of these patterns are illustrated in column 3 of the figure.) In the more typical case where the strips of positive and negative samples in row 3 of the figure show partial overlap (as in column 2 of the figure), the maximum value of *F*_0_(*x*) − *F*_1_(*x*) will be less than 1, but the closer this maximum is to 1, the greater is the ability of qPCR-based concentrations to discern the difference between the two groups. In summary, the more clearly the actual positive and negative samples are separated along the qPCR-based concentration axis, the higher the peak of the discernibility function will be, with a peak height of 1 indicating that the two groups can be distinguished without error.

Each type of plot in rows 2–7 is useful for giving a visual sense of the degree to which actual positive and negative samples separate along the qPCR-based concentration axis. Empirical density functions and empirical survival functions, however, suffer from a basic incompatibility with data sets that contain a non-negligible proportion of left-censored data, as will usually be the case for real monitoring data. The discernibility function is particularly useful for suggesting the particular qPCR-based concentration that permits maximum differentiation between actual positive and negative samples. Note that the greater the degree of separation between actual positives and negatives in the horizontal strip charts in row 3, the clearer will be the separation between *F*_0_(*x*) and *F*_1_(*x*) and between *S*_0_(*x*) and *S*_1_(*x*), the higher *F*_0_(*x*) will be when *F*_1_(*x*) first increases noticeably above 0, the lower *S*_0_(*x*) will be when *S*_1_(*x*) first decreases noticeably below 1, and the closer the maximum value of *F*_0_(*x*) − *F*_1_(*x*) will be to 1.

Fig. A.2 shows plots of *F*_0_(*x*) versus *F*_1_(*x*) (top row) and of *S*_1_(*x*) versus *S*_0_(*x*) (bottom row) for the three sets of data. Comparing the plots for the three data sets is useful for grasping the connection between the shape of these somewhat non-intuitive types of plot and the degree to which qPCR-based concentrations permit one to distinguish between actual positive and negative samples. They also nicely illustrate the usefulness of AUC (Area Under the Curve, calculated by numerically integrating the *F*_0_(*x*)-versus-*F*_1_(*x*) curve) as a numerical diagnostic index.

**Figure A.2.**
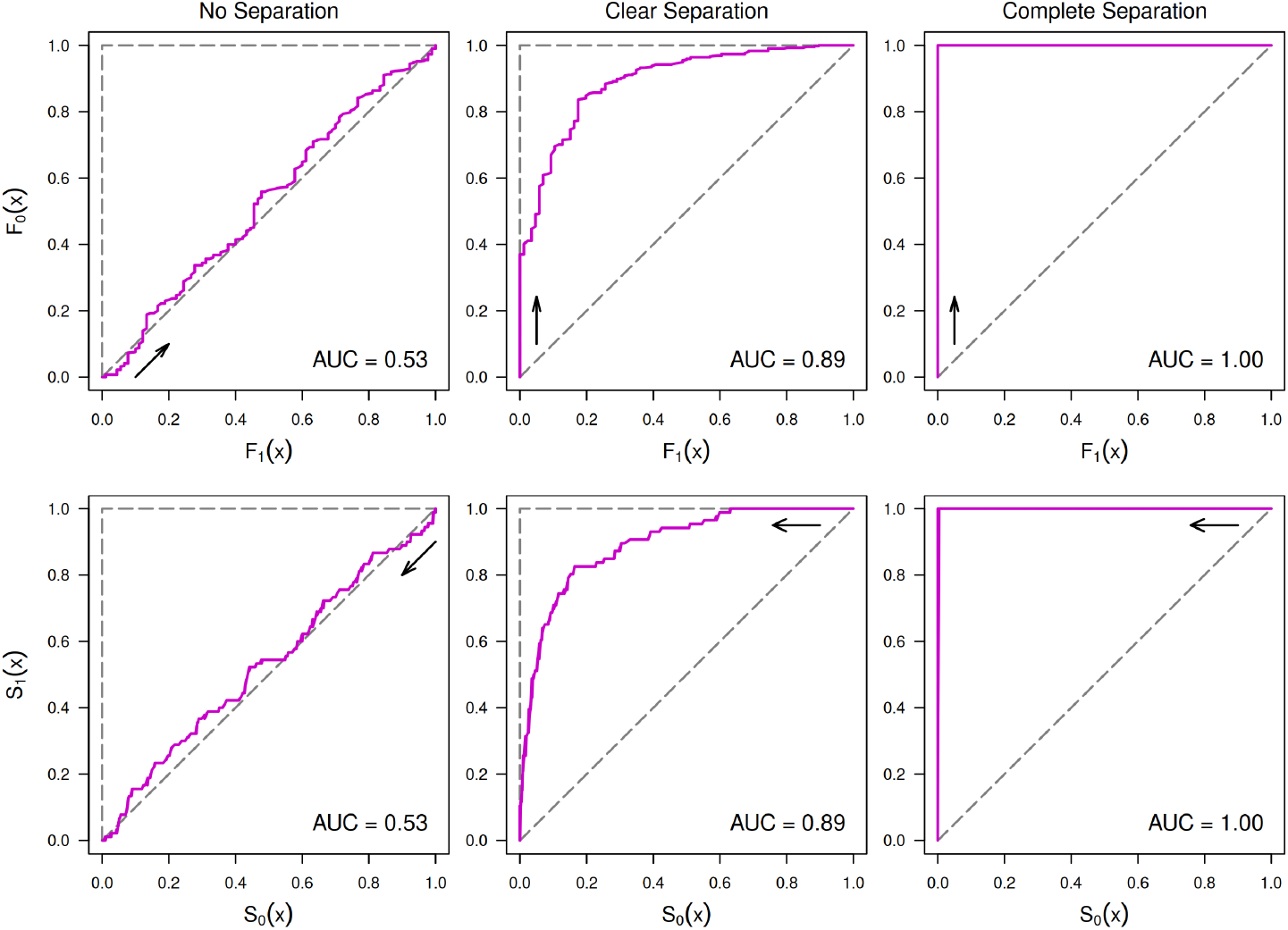
Plots of the relationship between empirical distribution functions *F*_0_(*x*) and *F*_1_(*x*) (top row) and between empirical survival functions *S*_1_(*x*) and *S*_0_(*x*) (bottom row) for the three data sets on which Fig. A.1 is based. Plots of *S*_1_(*x*) versus *S*_0_(*x*) are often called ROC (Receiver Operating Characteristic) plots. AUC denotes the Area Under the Curve. Arrows indicate direction of increasing *x*.

Plots of *S*_1_(*x*) versus *S*_0_(*x*) are often called ROC (Receiver Operating Characteristic) curves, a term that we feel obscures the true nature of the plot and that we agree is “not very enlightening” (Russell and Norvig, 2003, p. 843). It is also important to note that, because empirical survival functions are not appropriate for data sets that include a non-negligible proportion of left-censored qPCR-based concentrations, plots of *S*_1_(*x*) versus *S*_0_(*x*) (i.e., standard ROC plots) likewise are not appropriate for such data sets.

But as the examples in Fig. A.2 show, essentially the same information is contained in plots of *F*_0_(*x*) versus *F*_1_(*x*), which are appropriate for such data sets. Indeed, for uncensored data, it is straightforward to show analytically that the AUC values are identical for *F*_0_(*x*)-versus-*F*_1_(*x*) and *S*_1_(*x*)-versus-*S*_0_(*x*) curves, as these examples illustrate numerically. To see this, note that the area under an *F*_0_(*x*)-versus-*F*_1_(*x*) curve is simply the Stieltjes integral (or Lebesgue-Stieltjes integral, in the modern theory) of *F*_0_(*x*) with respect to *F*_1_(*x*) and can therefore be written as

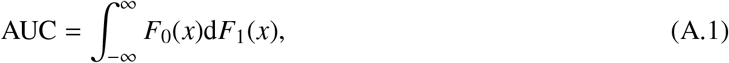

where distribution functions *F*_0_(*x*) and *F*_1_(*x*) may be continuous (e.g., theoretical distribution functions for continuous random variables) or step functions (e.g., EDFs). Integrating by parts, and noting that lim_*x*→∞_ *F*_0_(*x*)*F*_1_(*x*) = 1 and lim_*x*→−∞_ *F*_0_(*x*)*F*_1_(*x*) = 0, we find that

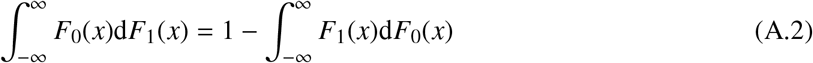

(Apostol, 1974: Chapter 7; Kolmogorov and Fomin, 1975: Section 36; Ross, 2013: Section 35). Recalling that *F*_*i*_(*x*) = 1 − *S*_*i*_(*x*) for survival function *S*_*i*_(*x*),

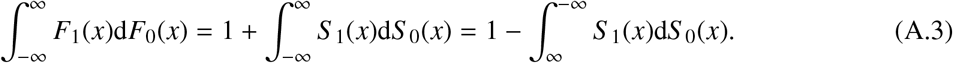

It now follows from Eqs. (A.1), (A.2), and (A.3) that

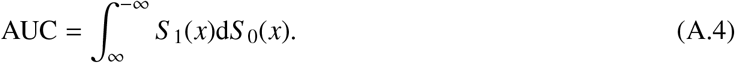

Note that the integral in Eq. (A.4) is calculated as *x* decreases, guaranteeing that *dS* 0(*x*) ≥ 0 and AUC ≥ 0.

### A.5 A Critique of Terminology for Performance Measures

As noted in Section 3.2 of the main text, five of the common synonyms for performance measures listed in Table 1 are misleading or simply incorrect. In each case, there is at least one alternative synonym that is logically valid and reasonably clear. The five misleading terms are “accuracy”, “precision”, “sensitivity”, “specificity”, and “selectivity”, whose use appears to be particularly popular in the medical literature dealing with performance assessment of diagnostic tests. We strongly recommend that these terms be avoided in science and engineering applications. The main reasons are as follows.

The percent of all decisions that are correct under a given decision rule is sometimes called the rule’s “accuracy”, but this is not what accuracy means in science. The concept of accuracy is commonly used in the physical sciences and applies to quantitative measurements (not the outcomes of binary decisions), is a relative rather than absolute concept, is not represented by a numerical value, and is defined qualitatively in terms of the sharper complementary concept of measurement error. The *International Vocabulary of Metrology* (BIPM et al., 2012), which is the main guidance document for measurement terminology in science, states that accuracy “is not a quantity and is not given a numerical quantity value. A measurement is said to be more accurate when it offers a smaller measurement error.” Measurement error, by contrast, is a quantitative concept and is expressed numerically. The term *percent error rate* (or simply *error rate*) for the percent of all decisions that are incorrect is consistent with this standard usage in science, and we recommend its use. If one prefers to emphasize the correct decisions instead of the incorrect ones, the term *percent correct decisions* (or simply *percent correct*) for the percent of all decisions that are correct is clear and valid. The term “accuracy” is avoided altogether in statistical theory, which instead employs the sharper complementary concept of bias.

The percent of putative positives, under a given decision rule, that are actual positives is sometimes called the rule’s “precision”, but here again, the term is both a traditional one from science that is being misused and one that is not used at all in statistical theory. As in the case of accuracy, precision in the physical sciences applies to quantitative measurements and, unless expressed inversely in terms of a statistical measure of variability, is a relative rather than quantitative concept. Still worse is the fact that in the present application, only the success rate in predicting positive cases is considered, thus ignoring the success rate in predicting negative cases, which cannot be inferred from the former. The commonly-used term *positive predictive value* and the clearer but less common term *true discovery rate* are far superior terms for the percent of putative positives that are actual positives. We also note that statistical theory does not employ the term “precision” but instead uses sharper complementary concepts based essentially on the notion of variance (e.g., standard deviation, standard error of the mean, 95% confidence interval).

The percent of actual positive samples that are correctly classified as positive is sometimes called the “sensitivity” of the decision rule. This usage, however, implies that a rule which blindly classifies all samples as positive, and is therefore completely insensitive to properties of the data, nevertheless has maximal (100%) sensitivity, which is absurd. It follows that the percent of actual positive samples that are correctly classified as positive does not measure sensitivity but merely the tendency to classify actual positive samples as positive, even if this classification is not based on properties of the data and therefore applies equally to actual negative samples. The basic problem is that the term “sensitivity” implies a response to a particular cue or stimulus (e.g., a specific DNA sequence), whereas the performance property being characterized (percent of actual positive samples that are correctly classified as positive) involves no such implication. A logically valid and reasonably clear term for this property is the *true-positive rate*, and we recommend its use.

A related issue is that the percent of actual negative samples which are correctly classified as negative is sometimes called the “specificity” or “selectivity” of the rule. But these usages imply that a rule which blindly classifies all samples as negative, and is therefore completely nonspecific and nonselective with respect to whatever properties distinguish actual negative samples from actual positive samples, nevertheless has maximal (100%) specificity and selectivity, which is absurd. As in the case of “sensitivity”, it follows that the percent of actual negative samples which are correctly classified as negative does not measure specificity or selectivity but merely the tendency to classify actual negative samples as negative, even if this classification is not based on properties of the data and therefore applies equally to positive samples. A logically valid and reasonably clear term for this property is the *true-negative rate*.

## Notes

### Competing Interest Statement

The authors have declared no competing interest.

### Summary of Updates

1. An inconsistency in formatting author addresses was fixed. 2. The name of the Michigan Network for Environmental Health and Technology was corrected.

